# Community-wide seed dispersal distances peak at low levels of specialisation in size-structured networks

**DOI:** 10.1101/2020.02.23.958454

**Authors:** Marjorie C. Sorensen, Matthias Schleuning, Isabel Donoso, Eike Lena Neuschulz, Thomas Mueller

## Abstract

Network approaches provide insight into the complex web of interspecific interactions that structure ecological communities. However, because data on the functional outcomes of ecological networks are very rarely available, the effect of network structure on ecosystem functions, such as seed dispersal, is largely unknown. Here, we develop a new approach that is able to link interaction networks to a trait-based seed-dispersal model to estimate community-wide seed dispersal distances. We simulated networks, using a niche model based on size-matching between plants and birds, that varied in the degree of niche partitioning. We found that community-wide dispersal distances were longest when networks had low degrees of niche partitioning. We further found that dispersal distances of plant species with small fruits peaked in models without niche partitioning, whereas dispersal distances of medium and large-fruited plants peaked at low degrees of niche partitioning. Our simulations demonstrate that the degree of niche partitioning between species is an important determinant of the ecological functions derived from ecological networks and that simulation approaches can provide new insights into the relationship between the structural and functional components of ecological networks.

## Introduction

During the last decade, studies of ecological networks have proliferated as a means to gain insight into the complex web of interactions between species (Heleno et al. 2014). Although ecological networks share some general properties such as an asymmetric distribution of links among species (Bascompte et al. 2006), how species partition their interaction partners varies widely across networks (Bascompte 2009). For instance, analysis of variation in network specialisation has shown that the degree of niche partitioning of pollination and seed-dispersal networks decreases with latitude (Schleuning et al. 2012), and that climate and human disturbance are important factors determining this variation in seed dispersal networks (Sebastian-Gonzalez et al. 2014). However, these studies only describe interaction frequencies between species in a community, while measures of the actual species contributions to the associated ecosystem functions are very rarely available across whole communities (but see Dennis and Westcott 2006, Rehm et al. 2019). Thus, the consequences of variability in network structure for community-wide ecosystem functions have not yet been quantified and investigated beyond conceptual considerations (Tylianakis et al. 2010, Blüthgen and Klein 2011).

An ecosystem function that is derived directly from interaction networks between plants and animals is animal-mediated seed dispersal. Seed dispersal away from the parent plant affects the dynamics, distribution, and long-term persistence of plant populations (Levin et al. 2003). Plant species with short seed dispersal ability may be unable to colonize new habitats, persist in fragmented landscapes, or respond to a changing climate (Turnbull et al. 2000, Neilson et al. 2005, Trakhtenbrot et al. 2005). An understanding of seed dispersal distances is, thus, essential for making predictions regarding future biodiversity change. Total dispersal kernels (TDKs; the frequency distribution of seed dispersal distances) offer an integrative measure of seed dispersal that account for the relative contribution of all major seed dispersers of a plant species (Jordano et al. 2007, Rogers et al. 2019, Nathan 2007). At the community-level, species-specific total dispersal kernels (TDK_plant_) can be integrated into a single community-wide total dispersal kernel denoted as TDK_community_ (analogous to TDK_system_ in Nathan 2007). This metric can be used to characterize differences in overall seed dispersal functions between communities and can serve as an estimate of community and ecosystem stability in response to global change (Loreau et al. 2003, Nathan 2007). Current possibilities for estimating seed dispersal simultaneously for several plant species include modelling (Schurr et al. 2009, Morales et al. 2013, Rehm et al. 2019, Rogers et al. 2019) and molecular approaches (Jordano et al. 2007, González-Varo et al. 2017), as well as approaches combining empirical data on frugivore movement and gut passage time (Holbrook and Smith 2000, Mueller et al. 2014). However, applying these methods to whole plant communities is a daunting task, because of the need to identify and quantify the relative contributions of numerous frugivore species to seed dispersal for every plant species in a community.

Functional traits are useful indicators of species’ ecological roles in ecosystems (Dehling et al. 2016) and may help overcome the challenge of quantifying the diverse contributions of frugivore species to seed dispersal. The matching between functional traits of consumer and resource species has a strong influence on network structure, due to its importance for determining which species interact preferentially with each other and how interaction partners are partitioned among species (Wheelwright 1985, Eklöf et al. 2013, Fründ et al. 2015, Dehling et al. 2016, Bender et al. 2018). For instance, large frugivores are more likely to consume large fruits, whereas small frugivores, constrained by a small gape width, are more likely to feed on small fruits (Cohen et al. 1993, Jordano et al. 2002, Eklöf et al. 2013, González-Castro et al. 2015, Bender et al. 2018). Consequently, size matching influences the degree of niche partitioning in plant frugivore networks (Dehling et al. 2014, Bender et al. 2018) and could have significant effects on seed dispersal because it directly affects which frugivores species will disperse which particular plant species.

Importantly, functional traits can also describe the ecological processes and functions that result from species interactions. For example, frugivore body size scales with gut passage time and movement distance, which means that large frugivores retain seeds longer and could carry them over greater distances than small frugivores (Robbins 1993, Yoshikawa et al. 2019). This results in longer-distance seed dispersal for plant species dispersed by large frugivores (Jordano et al. 2007, Wotton and Kelly 2012, Costa-Pereira et al. 2018). Past studies have estimated that a 100-fold increase in seed mass may result in a 4.5-fold increase in seed dispersal distance (Seidler and Plotkin 2006, Thomson et al. 2011). We suggest that the existing knowledge on how functional traits such as body size determine interactions between plants and frugivores, frugivore movement, and seed retention time, could help bridge the prevailing gap between network structure and ecological function.

Here, we propose a new approach that links interaction networks between plants and avian frugivores with a trait-based seed dispersal model to estimate dispersal kernels across whole plant communities. In order to examine how network specialisation is associated with seed dispersal distances at both the (1) community-wide (TDK_community_) and (2) individual plant species (TDK_plant_) level, we simulated networks with varying degrees of niche partitioning, using a niche model approach (Fründ et al. 2015, Donoso et al. 2017), while maintaining all other community parameters constant. We hypothesised that TDK_community_, and the majority of plant species TDKs, would be shorter in highly specialized networks because niche partitioning should result in the largest seed dispersers feeding only on a few plant species, contributing little to dispersal of the whole plant community.

## Material and methods

### Methods summary

1. *First, we simulated interaction networks that varied in the degree of niche partitioning and spanned a wide range of network specialisation using a niche model based on size-matching between plants and avian dispersers* (Fründ et al. 2015, Donoso et al. 2017); *Fig. 1a, b*).
2. *Second, we developed a trait-based seed dispersal model using allometric scaling relationships between avian body size, gut passage time, and flight speed (following Schurr et al. 2009) to estimate the seed dispersal distances provided by avian dispersers for each interaction in the simulated networks (Fig. 1c and Table 1 for an overview of model parameters*).
3. *Third, we combined the information from the simulated networks with the trait-based seed dispersal model to estimate dispersal distances for every plant species (TDK_plant_, Fig. 1d) and community-wide dispersal distances (TDK_community_) of each network*.
4. *Fourth, we conducted a global and a local sensitivity analyses on the parameters of the seed-dispersal model to test the robustness of our simulation model*.

**Figure 1.**
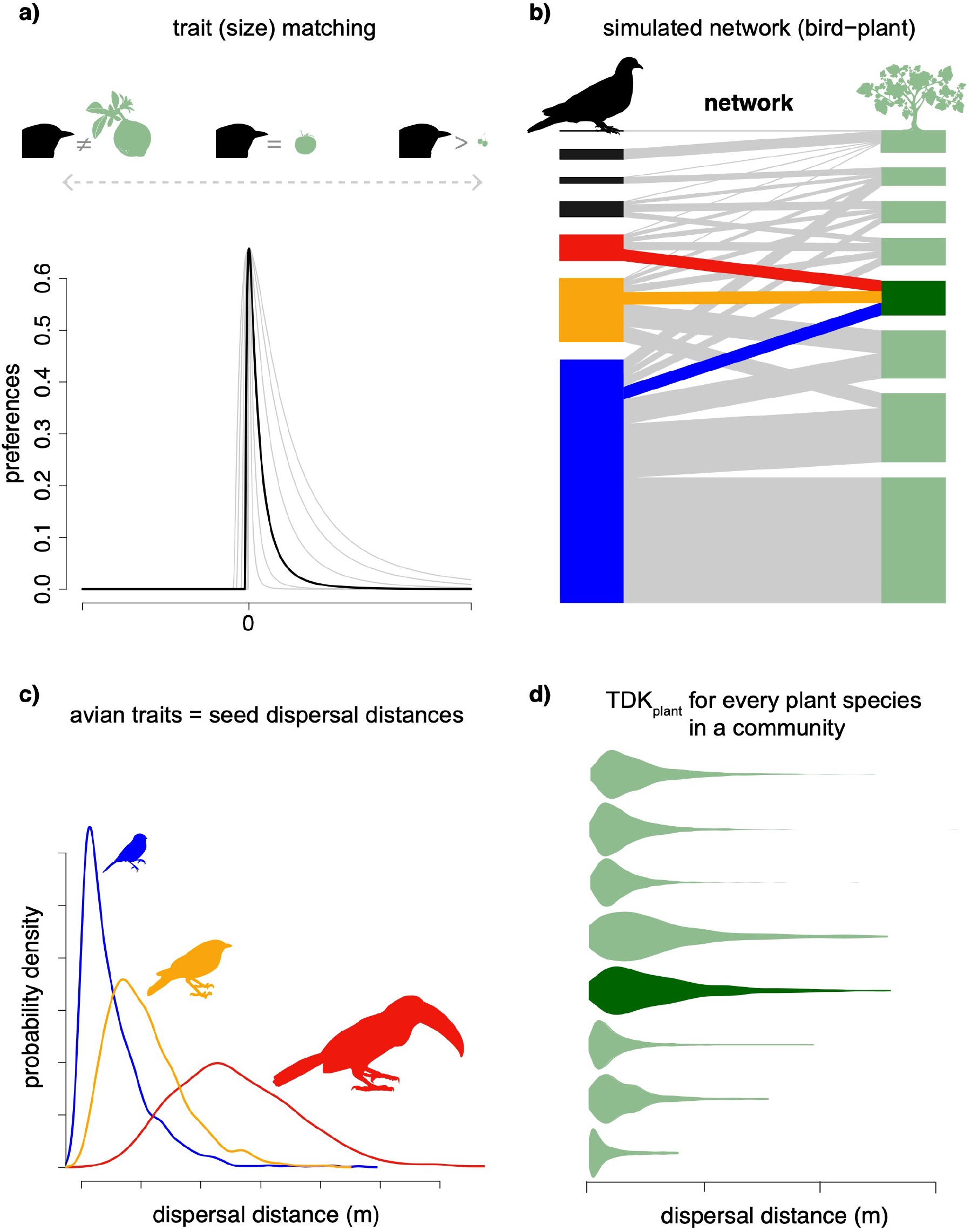
Estimating community seed dispersal (TDK_community_) with trait-based models. The proposed approach varies a) size matching between plant and frugivore species (derived from a right-skewed niche shape of trait matching as a function of trait distances between species; light grey lines indicate wider and narrower skewed niche shapes) to produce b) interaction networks with different degrees of niche partitioning (here, we display a single network with moderate specialisation). Models of interaction networks are combined with c) a trait-based model of frugivore movement to estimate dispersal kernels provided by each frugivore species to estimate d) dispersal distances for every plant species in a community (TDK_plant_). Colours indicate different frugivore species (blue = avian frugivore with small body size, orange = medium body size, red = large body size), and plant species (dark green = the plant species for which the method is illustrated, light green = all other plant species in the community).

**Table 1:** Summary information on model parameters included in the global sensitivity analysis.

| parameter | description | range |
| --- | --- | --- |
| gut passage time: |  |  |
| $GPT^{\text{exp}}$ | exponent of the GPT equation (3) | 0.39–0.62 |
| $GPT^{\text{var}}$ | variance of the GPT gamma distribution, $s^2$ in equation (5) and (6) | 2613–931509 |
| bird movement: |  |  |
| $FS^{\text{exp}}$ | exponent of the FS equation (4) | 0.13–0.21 |
| $FS^{\text{sd}}$ | standard deviation of the FS gaussian distribution | 0–4.7 |
| $CorrFactor$ | $fc$ in equation (7) | 0.001–0.004 |

#### 1. Simulating interaction networks with different degrees of niche partitioning

We used a simulation approach to build networks along the full gradient of specialisation, representing different degrees of niche partitioning (Donoso et al. 2017). This simulation approach allowed network specialisation to vary while maintaining all other community parameters, such as the number of plant and disperser species, constant. Simulations of size-structured interaction networks were based on trait distributions of avian body size and fruit volume in a theoretical community comprising 50 plant and 50 bird species. We focused on avian seed dispersers because birds are responsible for the majority of fruit removal (e.g., according to Jordano et al. 2007: 75 % birds, 15 % mammals).

For the simulations, species trait values were drawn from an idealized lognormal distribution with equidistant quantiles (Donoso et al. 2017), and the mean and standard deviation of body mass and fruit volume were defined by a large empirical data set of bird and fruit traits (n = 173 bird and 213 plant species; Bender et al. 2018). The total interaction frequencies of birds and plants were defined as a function of avian size and fruit volume (according to Donoso et al. 2017, see Supplementary material Appendix 1 for details), because smaller fruited plants are more abundant than larger fruited plants and smaller frugivores are more abundant than larger frugivores (Cotgreave 1993, Moles et al. 2005). In the simulations, the total number of interactions per bird species was kept fixed. Total interaction frequencies of plant species could vary among different model runs because they depended on the simulated bird preferences (Donoso et al. 2017).

According to the quantitative niche model used for simulating the networks (Donoso et al. 2017), we determined the preference of a bird species for a plant species as a function of the pairwise difference in trait values between bird body mass and plant fruit volume (Fig. 1a). Size matching in seed-dispersal networks is primarily driven by fruit size and avian gape width (e.g. Dehling et al. 2016). Since avian gape width and body mass are closely correlated (Moran and Catterall 2010), we chose to use body mass as this trait was also used for the simulations of seed dispersal distances (see below). We used a right-skewed niche shape to account for the fact that negative mismatches in trait values (bird < fruit) render interactions impossible (‘forbidden links’; Jordano et al. 2002), whereas positive size matching (bird > fruit) makes interactions less likely (Dehling et al. 2016). We modelled that birds choose among plants with a probability proportional to the product of preference and the total number of plant species interactions. By varying the breadth of bird foraging preferences, we were able to simulate different degrees of niche partitioning. In total, we simulated 116 networks, including a scenario without foraging preferences. We determined network specialisation for each simulated network by calculating the degree of complementary specialisation (*H_2_′*), a measure of niche partitioning ranging between 0 and 1, using the R package *bipartite* v. 2.11 (Dormann et al. 2009). For additional technical details on simulating interaction networks see the Supplementary material Appendix 1.

#### 2. Trait-based seed dispersal distance model

To estimate the seed dispersal distance resulting from each plant-bird interaction in the simulated networks, we developed a trait-based seed dispersal distance model. The two main components determining seed dispersal distances provided by frugivorous birds were: 1) gut passage times, and 2) movement distances (Westcott and Graham 2000, Jordano et al. 2007).

### Gut passage time

Since larger birds have longer gastrointestinal tracts, avian body mass and gut passage time (GPT) generally follow an allometric relationship (Robbins 1993). To build on the allometric relationship by Robbins 1993 and to develop an equation specifically for frugivorous birds foraging in natural environments, we collected GPT estimates from the literature. We only included studies that fed natural fruit to birds and excluded studies using artificial seeds or fruits, or marker dyes (see Supplementary material Appendix 1 Table A1 for included studies). We found a strong positive relationship between body mass (BM) and gut passage time (GPT; r^2^ = 0.69 p < 0.001; Supplementary material Appendix 3 Fig. A1). We used ordinary least squares (OLS) to estimate the steepness of the scaling relationship (Kilmer and Rodríguez 2016), resulting in the equation:

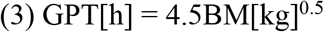

where GPT[h] is gut passage time, and BM[kg] is body mass. We focus on dispersal events resulting from endozoochory via defecation, although seeds can also be dispersed by other means such as epizoochory (Sorensen 1986) and regurgitation (Kays et al. 2011). We also did not include fruit size effects on GPT since observed patterns are inconsistent across studies, and include negative and positive relationships between seed size and GPT (Fukui 2003, Lenz et al. 2011, Wilson and Downs 2012).

### Movement distance

Body size generally scales positively with movement distance across several animal taxa including birds (Turner et al. 1969, Minns 1995, Carbone et al. 2005, Ottaviani et al. 2006); however, there is no reliable information on the general relationship between body size and home range size for bird species as birds often make movements beyond their home range (Lenz et al. 2011). We thus used the allometric equation between bird body mass and flight speed (FS) developed by (Tucker 1974) as a metric of foraging distance (Schurr et al. 2009, Tsoar et al. 2011, Viana et al. 2016):

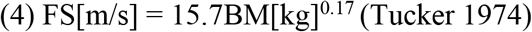

where FS is flight speed (in no-wind conditions), and BM is body mass. Equation 4 theoretically derives flight speed from avian aerodynamic measures collected during wind tunnel experiments (Tucker 1974).

### Combining gut passage time and movement distance

We used the allometric relationships between bird body mass - GPT (3), and bird body mass - movement distance (4), to parameterize a trait-based seed dispersal model, building on earlier studies that used similar approaches for individual species (Schurr et al. 2009, Tsoar et al. 2011, Viana et al. 2016).

For every interaction between a bird and plant, we followed the process of fruit consumption and passage through the gut until elimination. First, gut passage time was drawn from a Gamma distribution (Guttal et al. 2011). We chose a Gamma distribution because it most closely matches the GPT data found in empirical studies (Guttal et al. 2011, Pires et al. 2017). The shape (*k*) and scale (*θ*) parameters of the Gamma distribution can be defined in terms of the empirical GPT mean 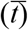 and variance (*s*^2^) as follows:

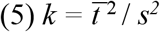

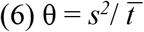

We used the allometric relationship (3) between body size and GPT to calculate the mean 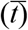. We selected a fixed variance which was set at the mean variance (*s*^2^) found in GPT studies collected during the literature search (Supplementary material Appendix 3 Table A1; (Pires et al. 2017)). Second, we selected the avian travel speed from a Gaussian distribution (Bruderer and Boldt 2008). We parameterized the Gaussian distribution using the mean flight speed calculated from allometric equation (4), and the average standard deviation of flight speeds reported in (Alerstam et al. 2007). We excluded birds larger than 1.5 kg from the standard deviation calculation because avian frugivores rarely exceed this size (Bender et al. 2018, Albrecht et al. 2018). Finally, we determined seed dispersal distance by multiplying GPT and FS.

Following Schurr et al. 2009, we calculated a calibration term of the simulated seed dispersal distances to account for the time frugivores spent not moving, and movements deviating from a straight line. To estimate the calibration term of absolute seed dispersal distances, we combined the GPT equation (equation 3 with hours converted to seconds) and flight speed equation (4) to produce the following:

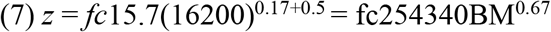

where *z* is seed dispersal distance (m) and BM is avian body mass (kg); *c* is a straightness factor which accounts for movements deviating from a straight line (*c* is 1 if movement occurs in a straight line); *f* is time allocated to movement as a constant fraction of the GPT (Schurr et al. 2009). We compared the independent expectation of the relationship between bird body mass and seed dispersal distance (equation 7) to the allometric equation derived from available empirical seed dispersal studies (*z* = 504BM^*0.48*^, Supplementary material Appendix 3 Table A2 Fig. S2). The calibration term (defined by the product of *f* and *c*), was calculated by computing the ratio between the allometric constant from equation 7 and that derived from empirical studies (504/254340). This resulted in a calibration term of *fc* = 0.002 which was applied to the simulated seed dispersal distance.

#### 3. Community-wide seed dispersal distance estimates

The model of seed dispersal was used to estimate the seed dispersal distance resulting from every plant-bird interaction in every simulated network (Fig. 1). We pooled the simulated seed dispersal distances for each individual plant species to create total dispersal kernels (TDK_plant_) for every plant species in each community (Fig. 1d). In order to estimate community-wide seed dispersal distances (TDK_community_), we calculated the median of the mean seed dispersal distance of all plant species in the respective network (each plant species was given equal weight when calculating the median). Similarly, we quantified community-wide long-distance seed dispersal (LDD) by taking the median of LDD events for each individual plant species. We defined LDD events as those beyond the 95^th^ percentile of the distribution of seed-dispersal distances. These calculations were repeated for each network along the full range of network specialisation.

Finally, we compared plants with different fruit sizes in order to investigate the association between network specialisation and TDK_plant_ for plant species with different sized fruits. Plants were grouped into small (bottom 25 % of species arranged by decreasing fruit volume), medium (middle 50 % of species arranged by decreasing fruit volume), and large (top 25 % of species arranged by decreasing fruit volume) fruits. All analyses and fitting of *loess* smoothing curves were conducted in R version 3.5.0 (R Core Development Team 2018; the *loess* smoothing parameter was equal to 0.2 for all figures).

#### 4. Sensitivity analysis

To estimate the influence of different model parameters on the seed dispersal distance model, we carried out a global sensitivity analysis. We used the Morris’s elementary effects method (Morris 1991) which estimates the relative rank of parameter importance while taking into account parameter interactions and is the most appropriate method for individual-based simulation models (Thiele et al. 2014). *μ*\* provides the order of importance for each factor with respect to the model output and can be considered as a proxy of the total sensitivity index (Supplementary material Appendix 5). We performed the sensitivity analysis on six model parameters, which were varied according to predefined ranges (Table 1). The sensitivity analysis was performed using the R package *sensitivity* version 1.16.0 (Pujol et al. 2015).

Based on the results of the global sensitivity analysis, we applied a local sensitivity analysis to test whether variation in the relevant parameters influenced our main findings. To this end, we selected the three most important model parameters for each of the mean and 95% quantile model outputs and evaluated the relationship between network specialisation (*H_2_′*) and TDK_community_ for both the maximum and minimum values of each of the important model parameters (see Table 1 for the range of variation in the model parameters).

## Results

### Community level

Community-wide seed dispersal distances (TDK_community_) varied systematically along the range of network specialisation. Networks with a low degree of niche partitioning showed longer community-wide seed dispersal distances than networks with no niche partitioning or high partitioning, resulting in a hump-shaped relationship between network specialisation and community seed dispersal distance (Fig. 2a). Mean community seed dispersal distances were longest at *H_2_′* = 0.11 (74 m) and shorter at both *H_2_′* = 0 (59 m) and *H_2_′* = 0.98 (46 m). Community-wide long-distance seed dispersal (LDD; 95 % quantile), as an alternative descriptor of TDK_community_, resulted in a similar hump-shaped relationship between network specialisation and LDD (Fig. 2b). Community-wide LDD events were longest when niche partitioning was low, *H_2_′* = 0.07 (173 m), and shorter at both, *H_2_′* = 0 (166 m) and *H_2_′* = 0.98 (65 m). However, LDD distance declined more rapidly between low and high degree of niche partitioning than mean seed dispersal distances (mean = 38 % decline between *H_2_′* = 0.11 and *H_2_′* = 0.98, LDD = 63 % decline between *H_2_′* = 0.07 and *H_2_′* = 0.98).

**Figure 2.**
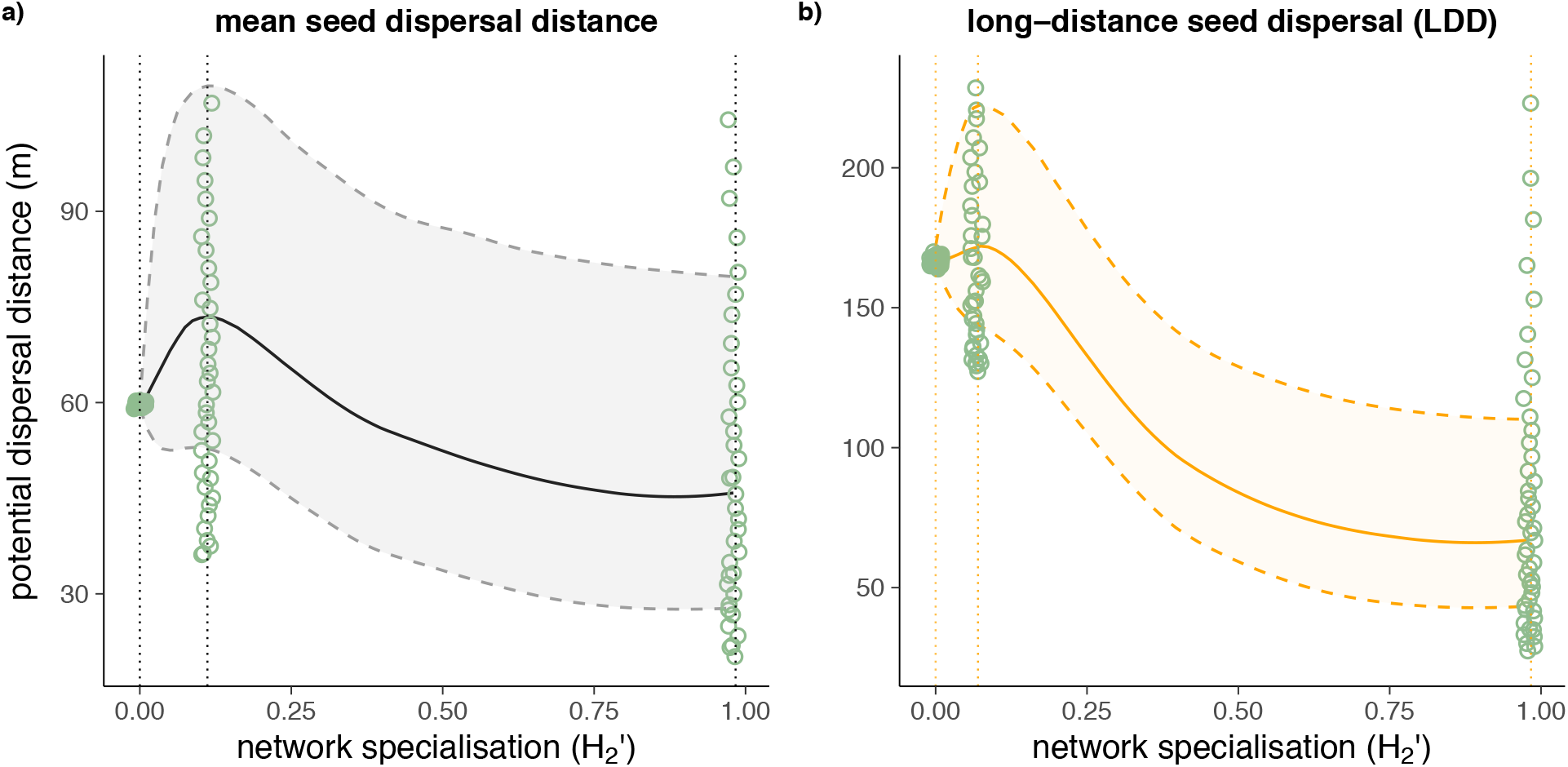
Relationship between network specialisation (*H_2_′*) and community-wide a) mean seed dispersal distances, and b) LDD (95 % seed dispersal distance quantile) for the overall community (representing two alternative descriptors of TDK_community_). Dotted lines intersect the x-axis at *H_2_′* = 0 (no niche partitioning), *H_2_′* = 0.1 (maximum community seed dispersal or LDD), and at *H_2_′* = 0.98 (complete niche partitioning). At each point where dotted lines intersect the x-axis (*H_2_′*: 0, 0.11 & 0.98) open green circles show the mean or 95 % quantile seed dispersal distances for all plant species that fall within the plotted range. Please note different scales of the y-axes.

### Plant species level

Seed dispersal distances of individual plant species (TDK_plant_) were associated with plant species traits (Fig. 3). Mean seed dispersal distances of plant species with the smallest fruits declined steadily from no to complete niche partitioning (67 % decline between no niche partitioning and complete niche partitioning; Fig. 3b). In contrast, mean seed dispersal distances of plant species with medium sized fruits were longest when specialisation was low, *H_2_′* = 0.11 (75 m), and shorter at the extremes of niche partitioning (no niche partitioning = 59 m; complete niche partitioning = 45 m; Fig. 3c). Mean seed dispersal distances of plant species with the largest fruits were also longest when niche partitioning was low (156 m) and shorter at the extremes of network specialisation (no niche partitioning = 59 m; complete niche partitioning = 126 m; Fig. 3d). Minimum seed dispersal distances of the largest fruited plant species occurred when networks had no niche partitioning; whereas, minimum seed dispersal distances for medium and small fruited plants occurred when niche partitioning was highest. Long-distance seed dispersal, as an alternative descriptor of TDK_plant_, followed the same pattern (Supplementary material Appendix 4 Fig. A4).

**Figure 3.**
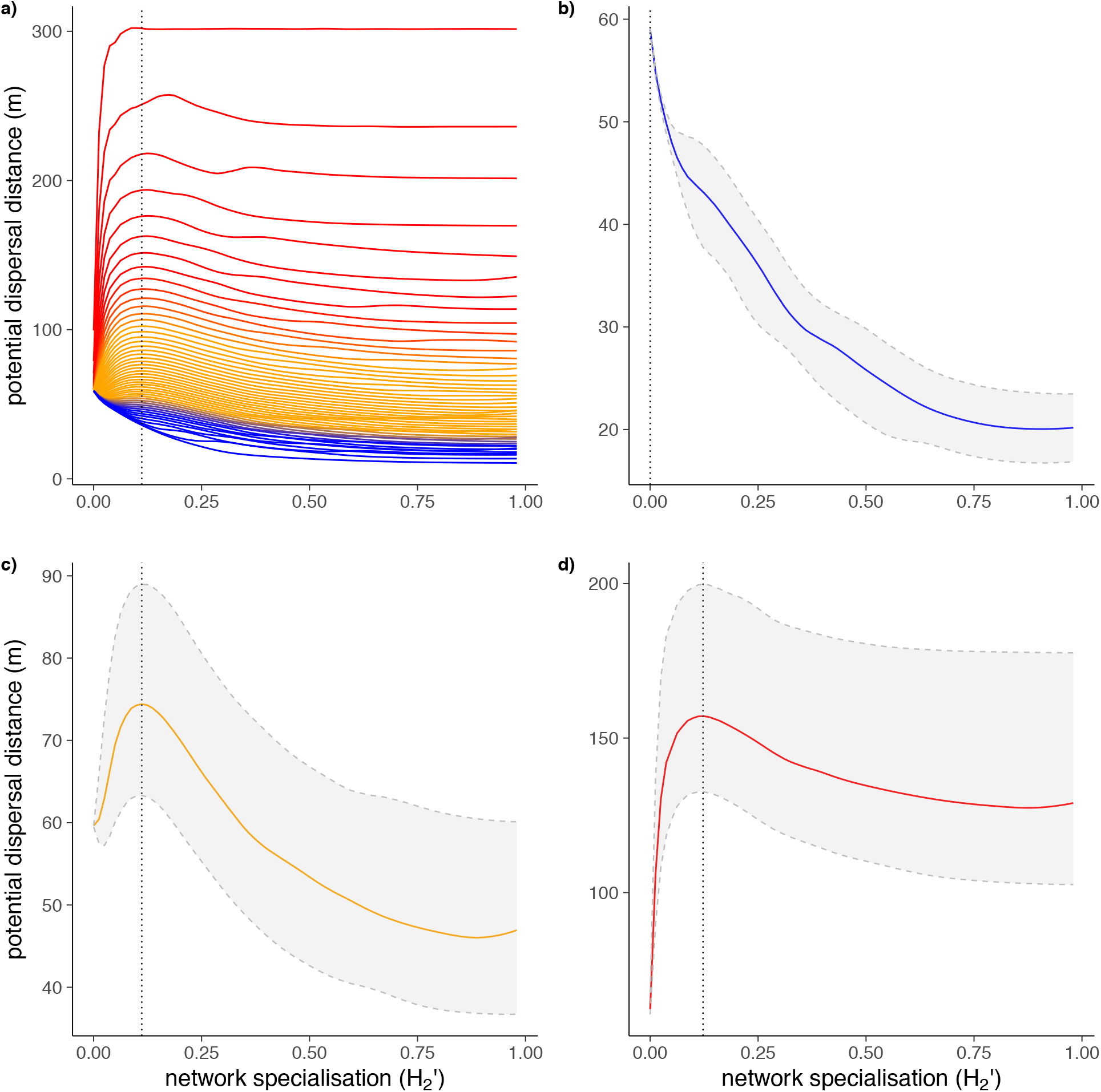
Relationship between network specialisation (*H_2_′*) and seed dispersal distances for plants (TDK_plant_) with different fruit sizes. The colour gradient ranges from blue (small fruited plant species, smallest 25 %) to orange (medium fruited plant species, middle 50 %) to red (large fruited plant species, largest 25 %). a) Mean seed dispersal distances for each plant species in the community. Median of mean seed dispersal distances b) for plant species with the smallest fruits, c) for plant species with medium sized fruits, and d) for plant species with the largest fruits. Dotted lines intersect the x-axis at maximum seed dispersal distances (community-wide: *H_2_′* = 0.1; small fruits: *H_2_′* = 0; medium fruits: *H_2_′* = 0.1; large fruits: *H_2_′* = 0.11). The light grey area represents the 75 % confidence intervals. Please note different scales of the y-axes.

### Sensitivity analysis

The sensitivity analyses showed that the simulation results were robust to variation in the parameter estimates. The most influential parameters for both the mean and 95% quantile of seed dispersal distances were *GPT*var, *CorrFactor*, and *GPT*^exp^, while the parameters of the FS equation were of little relevance (Fig. 4). Although absolute seed dispersal distances varied in the local sensitivity analysis, the relationship between network specialisation and community-wide seed dispersal distance (TDK_community_) remained qualitatively consistent, varying *GPT*var, *CorrFactor*, and *GPT*^exp^ (Supplementary material Appendix 5 Fig. A5 – A7).

**Figure 4.**
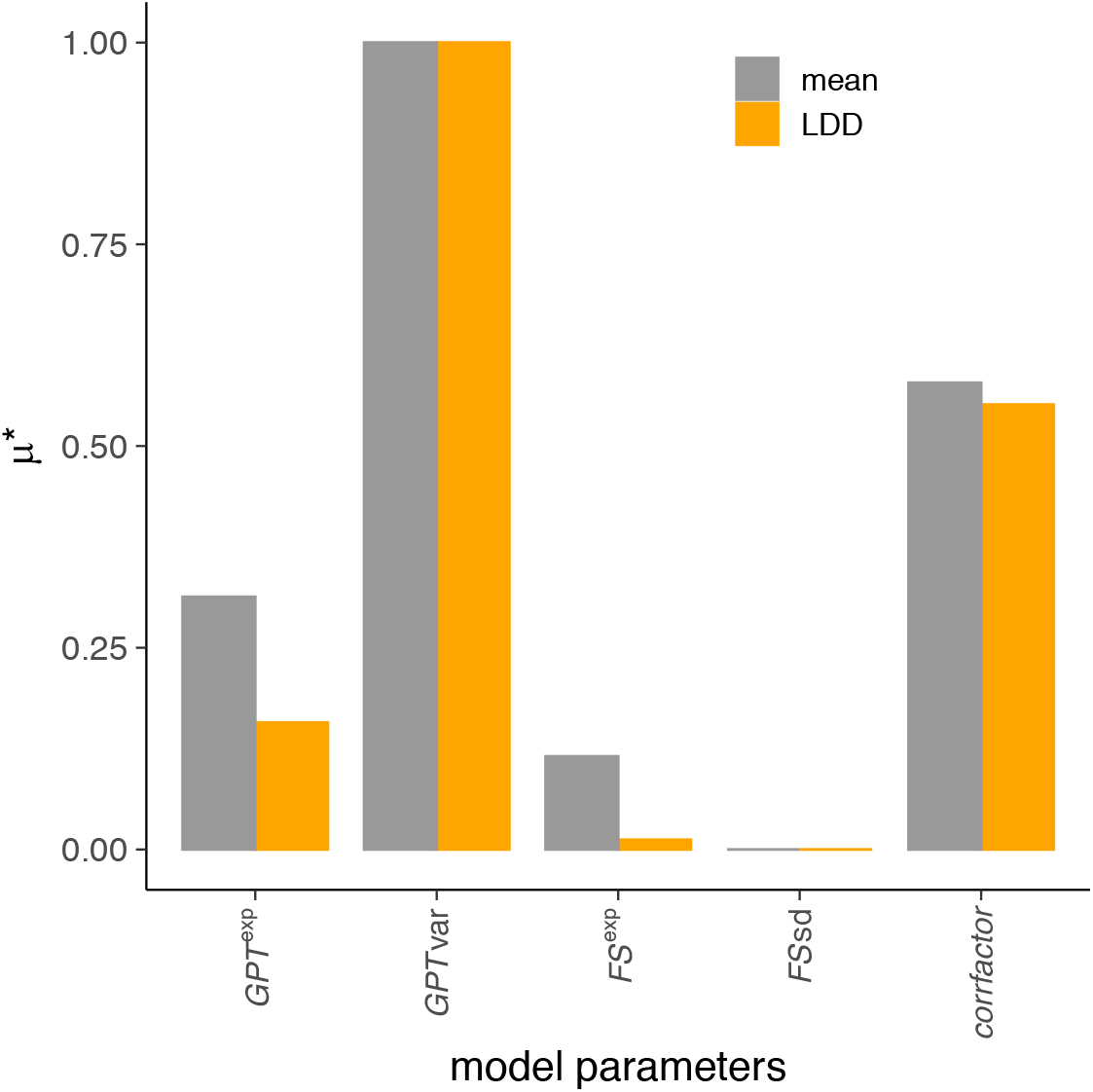
Results from the sensitivity analysis for two different descriptors of TDK_community_: mean seed dispersal, and LDD (95 % quantile of seed dispersal distances). Bars show the μ*values ranking the relative influence of each model parameter on the results for both descriptors (grey = mean seed dispersal distance; orange = LDD). See Table 1 for a full description of each model parameter under consideration.

## Discussion

We propose a new approach to link interaction networks with a trait-based seed dispersal model to estimate avian seed dispersal distances of plant communities. We found support for the hypothesis that network specialisation is systematically associated with the total dispersal kernels of plant communities (TDK_community_). Specifically, we found that the mean and LDD of community-wide seed dispersal distance, as two alternative descriptors of TDK_community_, were longest when niche partitioning between bird and plant species was low, and shorter in scenarios of complete and no niche partitioning. This hump-shaped relationship between seed dispersal and network specialisation was driven by changes in the relative contribution of birds to the seed dispersal of medium and large-fruited plants at different scenarios of niche partitioning. These results suggest that low niche partitioning between plants and avian frugivores maximizes community-wide seed-dispersal functions.

Our simulations demonstrate that variation in the degree of niche partitioning is associated with ecosystem functioning via effects on TDK_community_. The observed hump-shaped relationship between seed dispersal distances and network specialisation results from different avian foraging preferences at different levels of niche partitioning. Reduced seed dispersal in networks with no niche partitioning (*H_2_′* = 0) may be explained by small and large bird species being able to feed on all plant species, leading to weak effects of large birds on community seed dispersal. At the other extreme, in networks with complete niche partitioning (*H_2_′* = 1), large birds interacted with only a few large-fruited plant species and, thus, had comparatively little effect on community-wide dispersal distances. The effects of large birds species were maximized at low niche partitioning (*H_2_′* = 0.11) because under these conditions large species were able to forage widely across the plant community, whereas small birds were restricted to small-fruited plants, due to morphological size constraints. We found that LDD events also peaked when niche partitioning was low (*H_2_′* = 0.07), and large birds were foraging most widely; however, LDD declined more rapidly than mean seed dispersal as network specialisation increased. This is consistent with empirical studies which have shown that long-distance dispersal usually results from fruit removal by the largest bird species (Wotton and Kelly 2012, Mueller et al. 2014, Costa-Pereira et al. 2018).

The results of our simulation study have implications for real-world communities which vary widely in the degree of niche partitioning between plants and avian frugivores. Empirical studies have shown that plant-frugivore networks vary in network specialisation (*H_2_′*) between 0.16 – 0.58 and exhibit this structural variability at both global and local scales (Schleuning et al. 2012, Quitián et al. 2018). Our results suggest that size-structured networks within this range of network specialisation may vary both in community and individual plant species seed dispersal distances. For example, networks with a low degree of niche partitioning, such as networks at forest edges (Menke et al. 2012) and at low latitudes (Schleuning et al. 2012), especially in the Afrotropics (Dugger et al. 2018), may provide longer community seed dispersal distances than the comparatively more specialised networks in forest interiors and at higher latitudes. However, our simulation study was based only on variation in the degree of size matching between species and kept community and trait composition constant. In addition to effects of size matching, variability in network structure is also driven by other factors such as the spatial and temporal fluctuations in resource availability and species abundances (Carnicer et al. 2009, Bender et al. 2017, Sebastian-Gonzalez et al. 2017). Nevertheless, our study demonstrates that variation in the degree of size matching between species alone can trigger substantial differences in seed dispersal.

The functional outcome of network structure that we have measured in terms of TDK_community_ may serve as an important measure of community and ecosystem stability in response to environmental change, as has been conceptually suggested in previous studies (Loreau et al. 2003, Nathan 2007). Previous studies seeking to understand the potential consequences of variability in network structure primarily investigated how network structure is related to community stability. For instance, highly generalised and connected networks are more resistant to secondary extinctions following species loss (Memmott et al. 2004, Okuyama and Holland 2008, Thébault and Fontaine 2010, Rohr et al. 2014), are less likely to disassemble (Sole and Montoya 2001, Dunne and Williams 2009), and are more resistant to species invasions (Post and Pimm 1983) than more specialised networks. This stability is likely due to an association between niche partitioning and functional redundancy, as similar species can fulfill similar functional roles and compensate for species loss in generalised networks (Blüthgen and Klein 2011). Here, we move beyond structural measures of community stability by estimating the functional outcome derived from species interactions in ecological networks. We show that a generalised structure of ecological networks leads to a higher degree of ecosystem functioning. We suggest that TDK_community_ could be used as a community-wide indicator for assessing the stability of ecosystems in response to global change. Insufficient dispersal distance constrains the adaptive capacity of species to changing climatic conditions, for example by reducing the speed of plant range shifts (Neilson et al. 2005, Trakhtenbrot et al. 2005) and plant persistence in fragmented landscapes (Turnbull et al. 2000).

We found that the association between network structure and seed dispersal was mediated by plant species traits. Seed dispersal distances of small-fruited plant species were longest when niche partitioning was completely absent since in this scenario the largest bird species feed equally across plant species. In contrast, seed dispersal distances of medium and large-fruited plant species were the drivers of the community-wide hump-shaped relationship between dispersal distance and network specialisation. At low degrees of niche partitioning, these plants received a higher proportion of fruit removal by large frugivores compared to small frugivores because asymmetric size matching renders interactions between large fruits and small birds impossible.

The association between plant species traits and niche partitioning could influence the spatial patterns of seed dispersal for different types of plant species. Since seed dispersal is the critical first step for competitive processes and subsequent recruitment (Nathan and Muller-Landau 2000, Rohr et al. 2014), spatial patterns of seed dispersal may have implications for long-term persistence and coexistence of plant species. Theoretical work has highlighted conspecific spatial clustering as a mechanism reducing competitive exclusion and promoting diversity (Hubbell 1986, Chave et al. 2002). Empirical work has demonstrated that between-species variability in seed dispersal distances leads to differences in spatial clustering of plants, such that species with shorter seed dispersal distances are more tightly clustered in space than species with longer seed dispersal distances (Seidler and Plotkin 2006). In our simulations, we found no variability between seed dispersal distances across plant species under a scenario that lacked niche partitioning (Fig. 2; *H_2_′* = 0); whereas, increasing niche partitioning resulted in higher variability between species-specific seed dispersal distances (Fig. 2; *H_2_′* = 0.1). This suggests that niche partitioning among avian frugivores may contribute to conspecific aggregation and coexistence of plant species.

The trait-based model proposed here combines simulated ecological networks and allometric relationships to estimate the functional effects of species interactions in complex ecological networks. We show that this trait-based approach can be used to estimate avian seed dispersal for whole plant communities. The simplicity of this approach allows the model to be broadly applicable to plant communities that are primarily dispersed by birds and for which trait information are available. However, other animal frugivores may also be important seed dispersers (e.g., Mello et al. 2011, Dugger et al. 2018), and our model would need to be developed further to also account for the contributions of other animal frugivores (but see Pires et al. 2017) for a similar approach for mammal seed dispersers). While the main goal of our analysis was not to estimate absolute seed dispersal distances, we stress that the estimated distances closely match those expected from the few available empirical studies (Jordano et al. 2007). The sensitivity analysis of our simulation model showed that our main finding, derived from the comparison of seed dispersal distances among differently structured networks, was qualitatively consistent along the full range of selected model parameters (see Supplementary material Appendix 5 Fig. A5-A7). Nevertheless, we acknowledge that our trait-based approach is not able to capture variability in avian behaviour, independent of variation in body size. For example, the model cannot account for species differences in habitat selection, responses to resource availability, or mating strategies, all of which may affect movement and seed deposition (e.g., Wenny and Levey 1998, Karubian and Durães 2009, Morales et al. 2013, Da Silveira et al. 2016). The collection of empirical movement data from a wide range of animal species would help to improve the seed dispersal distance estimates of trait-based models and could cover additional species traits and more aspects of animal behaviour.

We conclude that our trait-based model provides a new means by which seed dispersal distances can be estimated for whole plant communities. Our simulation study demonstrates that variability in how species interact in ecological communities is relevant for determining the ecological functions derived from ecological networks. These findings show the relevance of better integration of structural and functional approaches in network ecology and should fuel more theoretical and empirical research on linking network structure and function.

## Acknowledgements

We are grateful to Jochen Fründ for suggestions and guidance on network simulations and to Javier Rodríguez-Pérez for input on the sensitivity analysis. MCS and ID were funded by the Alexander von Humboldt Foundation. TM was funded by the Robert Bosch Foundation. ELN and MS obtained funding by the German Research Foundation (DFG) in the framework of the Research Bundle 823–825 “Platform for Biodiversity and Ecosystem Monitoring and Research in South Ecuador” (PAK 825/1) and the Research Unit FOR2730 “Environmental changes in biodiversity hotspot ecosystems of South Ecuador: RESPonse and feedback effECTs”.

