## Supplementary material for "Community-wide seed dispersal distances peak at low levels of specialisation in size-structured networks"

**Contents**

**Appendix 1**: Additional details about the interaction network simulation approach

**Appendix 2**: Source code for the mechanistic trait-based seed dispersal model in the R language for statistical computing

**Appendix 3**: Additional details about the trait-based seed dispersal model and allometric equations used

Table A1: Summary of feeding trial studies for the relationship between avian frugivore body mass and gut passage time

Table A2: Summary of field-based empirical studies for the relationship between avian frugivore body mass and seed dispersal distances

Figure A1: Relationship between body mass and mean gut passage time using data extracted from empirical feeding trials for frugivorous birds

Figure A2: Relationship between body mass and max seed dispersal distance using data extracted from empirical feeding trials for frugivorous birds

Figure A3: Relationship between body mass and mean dispersal distance using data extracted from empirical studies of seed dispersal by frugivorous birds

**Appendix 4**: Long-distance seed dispersal (LDD) for small, medium, and large fruited plant species

Figure A4: Small, medium, large fruited plant species long-distance seed dispersal

**Appendix 5**: Sensitivity analysis

Table A3: Model parameters included in the sensitivity analysis

Figure A5: Gut passage time exponent (GPTexp) local sensitivity analysis

Figure A6: Gut passage time variance (GPTvar) local sensitivity analysis

Figure A7: Correction factor (corrfactor) local sensitivity analysis

**Appendix 1: Additional details about the interaction network simulation approach**

In the network simulations, total interaction frequencies of plants took differences in plant species abundance into account. We assumed a negative relationship between fruit size and interaction frequency (Donoso *et al*. 2017; González-Castro *et al*. 2015; Moles *et al*. 2005):

(1) $f_i = 1/x_i$

where $x_i$ represents the fruit volume value for plant *i*, and $f_i$ represents the expected total interaction frequency (Donoso *et al.* 2017).

Similarly, total interaction frequencies of bird species took differences in bird abundance into account. We assumed a negative relationship between body mass and abundance (Cotgreave 1993; González-Castro *et al*. 2015); in this case, we assumed undercompensation (i.e. interaction frequency decreases less rapidly than bird size increases) as large birds tend to consume more fruits per individual (García *et al.* 2014).

(2) $g_j = (1/y_j) + \beta$

where $y_j$ is the bird size value for bird *j*, $g_j$ is the expected total bird interaction frequency, and $\beta$ being an undercompensation parameter, set to 10 % of the maximum value of 1/y. Donoso *et al*. 2017 found that results were robust to variation in the value of $\beta$.

**Appendix 2: Source code for the mechanistic trait-based seed dispersal model in the R language for statistical computing**

Explanatory comments (#)

```
#------------trait-based seed dispersal distance model----------------#


#The dispsimulation function generates estimated seed dispersal distances for plant-
 bird interactions. This function takes as input an object with disperser body mass
 (kg) for each interaction event in a network.

#Nbird = number of bird species in the community
#obsperbird = number of interaction events for each bird species

  dispsimulation <- function (x) {
  dispdist <- rep(NA, obsperbird * Nbird)
  for(i in 1:nrow(x)) {
#a mean GPT is selected from the allometric equation derived from empirical data
 presented in this study. [i,3] indicates the column where bird body mass is located,
 this may not fit with other data structures
    meanGPThour <- 4.5*x[i,1]^0.5

 #convert GPT to seconds (since speed is in m/s)
    meanGPT <- meanGPThour*3600

 #calculate the shape and scale parameters for the gamma distribution using meanGPT
      and variance (we chose variance = 100241, since this was the average GPT
      variance calculated across 11 empirical studies in which variance was reported;
      see Table S2)
    scalevalue <- 75311 /meanGPT
    shapevalue <- meanGPT^2/ 75311

 #select a GPT value for this particular interaction from the GPT gamma distribution
    GPT <- rgamma(1, shape = shapevalue, scale = scalevalue)

 #then select a mean flight speed (calculated used the allometric equation presented
 in Alerstam et al. 2007)
    meanspeed <- 15.7*x[i,1]^0.17

 #select a flight speed value for this particular interaction using meanspeed and
 2.078 to parameterize rnorm. 2.078 is the flight speed sd average
 reported in Alerstam et al. 2007 for those species with body mass lower than 1.77 kg
 (which is the largest bird species across our 7 Andean communities)
    speed <- rnorm(1, meanspeed, 2.078)

 #calculate the max distance travelled (if flying straight without stopping) given the
 selected GPT.
    max_distance <- speed*GPT

 #correction factor which accounts for birds resting/not always moving in a straight
 line.
    distance <- 0.002 * max_distance
```

```
 # NOTE! there may be a few cases where the speed value -selected from rnorm- could
 have a negative value.
 # For these few cases, the negative seed dispersal distance is replaced with NA.
    if (distance < 0){
      distance<-NA
    }
       dispdist[i] <- distance
  }
  return(dispdist)
}
```

### Appendix 3

**Table A1.** Summary of feeding trial studies for the relationship between avian frugivore **body mass** and **gut passage time.**

| Species | Body mass (g) | Mean retention time (min) | Std. deviation | Source |
|---|---|---|---|---|
| *Acanthagenys refogularis* | 44 | 40.6 | 12.5 | Murphy *et al*. 1993 |
| *Acridotheres cristatellus* | 123 | 18.4 | NA | Shi *et al*. 2015 |
| *Alophoixus pallidus* | 42.8 | 44 | 11 | Khamcha *et al*. 2014 |
| *Arizelocichla milanjensis* | 54 | 44 | NA | Lehouck *et al*. 2009, personal communication |
| *Bombycilla cedrorum* | 32 | 26.7 | 27.38 | Ramirez & Ornelas 2009 |
| *Bycanistes bucinator* | 635 | 64 | 29 | Lenz *et al*. 2011, personal communication |
| *Ceratogymna atrata* | 1431 | 248.4 | 124.6 | Holbrook & Smith 2000 |
| *Ceratogymna cylindricus* | 1038 | 218.4 | 95.2 | Holbrook & Smith 2000 |
| *Dicaeum hirundinaceum* | 9 | 13.7 | 6.6 | Murphy *et al*. 1993 |
| *Grantiella picta* | 20.7 | 24.4 | 9.77 | Barea 2008 |
| *Hemiphaga novaeseelandiae* | 650 | 120 | 39.1 | Wotton *et al*. 2008; Wotton *et al*. 2012 |
| *Hypsipetes amaurotis* | 78.7 | 20.8 | NA | Fukui 2003 |
| *Megalaima asiatica* | 90.5 | 26.9 | NA | Shi *et al*. 2015 |
| *Megalaima nuchalis* | 87.7 | 26.9 | NA | Chang *et al*. 2012 |
| *Mionectes oleagineus* | 11.5 | 15.7 | NA | Westcott & Graham 2000 |
| *Musophaga johnstoni* | 250 | 69.6 | 17.6 | Sun *et al*. 1997 |
| *Myadestes melanops* | 32.1 | 24.5 | NA | Murray 1988 |
| *Nestor notabilis* | 870 | 140.4 | NA | Young *et al*. 2012 |
| *Notiomystis cincta* | 35 | 13.5 | NA | Trass 2000 |
| *Onychognathus morio* | 135 | 35 | NA | Mokotjomela *et al*. 2015 |
| *Onychognathus tristramii* | 120 | 135.1 | NA | Spiegel & Nathan 2007 |
| *Penelope obscura* | 1770 | 346 | NA | Guix & Ruiz 1997 |
| *Phainoptila melanoxantha* | 56 | 17.5 | NA | Murray 1988 |
| *Phyllastrephus placidus* | 34.5 | 80.36 | NA | V. Lehouck, personal communication |
| *Prosthemadera novaeseelandiae* | 105 | 37 | NA | O'Connor 2006 |
| *Pycnonotus aurigaster* | 44.4 | 22.6 | NA | Shi *et al*. 2015 |
| *Pycnonotus jocosus* | 27.4 | 24 | NA | Shi *et al*. 2015 |
| *Pycnonotus melanicterus* | 28.9 | 35 | 8 | Khamcha *et al*. 2014 |
| *Pycnonotus xanthopygos* | 40 | 34.7 | NA | Spiegel & Nathan 2007 |
| *Semnornis frantzii* | 57.3 | 26.6 | NA | Murray 1988 |
| *Sturnus vulgaris* | 71 | 42.3 | 16.5 | LaFleur *et al*. 2009; Karasov & Levey 1990 |
| *Tauraco corythaix* | 300 | 110.4 | NA | Mokotjomela *et al*. 2015 |
| *Tauraco hartlaubi* | 235 | 42.9 | NA | Lehouck *et al*. 2009, personal communication |
| *Turdus helleri* | 66 | 45.73 | NA | Lehouck *et al*. 2009, personal communication |
| *Turdus merula* | 100 | 39.35 | 68.3 | Morales *et al*. 2013 |
| *Turdus migratorius* | 79 | 48 | NA | Karasov & Levey 1990 |
| *Zosterops lateralis* | 11 | 24.75 | 30.45 | French 1996; Stanley & Lill 2002 |

We developed an allometric equation specific to frugivores. We only included studies that fed natural fruit to birds and excluded studies using artificial seeds or fruits, or marker dyes. We used the search strings "seed or fruit + gut + retention or passage". For some studies GPT medians were reported instead of means, if means could not be attained via author personal communication or digitisation from presented plots, the study was not included in our analysis. The allometric relationship between body mass and GPT presented by Robbins 1993 included data on 21 bird species across all diet types (including studies using liquid and marker dye to measure GPT). Only 4 of the 21 species were fed fruits. The 37 included species are widely distributed across the weight range of frugivore species found in the seven Andean communities. Generally, standard errors were reported instead of standard deviations; however, if standard errors and sample sizes were both reported we converted standard error to standard deviation.

**Table A2.** Summary of field-based empirical studies for the relationship between avian frugivore **body mass** and **seed dispersal distances**.

| Species | Body mass (g) | Mean dispersal distance (m) | Max dispersal distance (m) | Source |
|---|---|---|---|---|
| *Bycanistes bucinator* | 635 | 528 | 14790 | Mueller *et al*. 2014 |
| *Ceratogymna atrata* | 1431 | 1521 | 6919 | Holbrook & Smith 2000 |
| *Ceratogymna cylindricus* | 1038 | 1537 | 4628 | Holbrook & Smith 2000 |
| *Corythaeola cristata* | 1000 | 240.5 | NA | Sun *et al*. 1997 |
| *Dicaeum hirundinaceum* | 9.25 | 103.67 | 500 | Ward & Paton 2007 |
| *Hemiphaga novaeseelandiae* | 650 | 84.7 | 1469 | Wotton & Kelly 2012; Wotton *et al*. 2008 |
| *Mionectes oleagineus* | 11.5 | 26.16 | 86 | Westcott & Graham 2000 |
| *Musophaga johnstoni* | 250 | 137.5 | NA | Sun *et al*. 1997 |
| *Myadestes melanops* | 31.5 | 84.7 | 364.7 | Murray 1988 |
| *Onychognathus tristramii* | 119 | 1168 | 4800 | Spiegel & Nathan 2007 |
| *Phainoptila melanoxantha* | 58 | 84.9 | 504.7 | Murray 1988 |
| *Prosthemadera novaeseelandiae* | 105 | 222.5 | NA | O'Connor 2006 |
| *Pycnonotus xanthopygos* | 40.5 | 303 | 900 | Spiegel & Nathan 2007 |
| *Semnornis frantzii* | 63.25 | 62.6 | 215 | Murray 1988 |
| *Turaco schuettii* | 250 | 149 | NA | Sun *et al*. 1997 |
| *Turdus merula* | 100 | 89.48 | 2220 | Breitbach *et al*. 2012 |

We used ordinary least squares (OLS) to fit an allometric equation between bird body mass and mean seed dispersal distance for empirical field-based studies (Table S1). This resulted in the following equation: $z = 504\text{BM[kg]}^{0.48}$, where $z$ is seed dispersal distance and BM is disperser species body mass. The ratio between the allometric constant from the independent expectation (equation 7 in the main text; 504/254340) and the allometric constant from empirical studies presented here was used to calculate the correction factor (0.002; accounting for movements deviation from a straight line and time not moving).

**Fig. A1**. Relationship between **body mass** and mean **gut passage time** using data extracted from empirical feeding trials for frugivorous birds (see detailed information about the studies in Table S2). Body mass is positively related to mean gut passage time ($r^2 = 0.69$, $p < 0.0001$, n=39). The grey shaded region indicates the confidence interval for the regression.

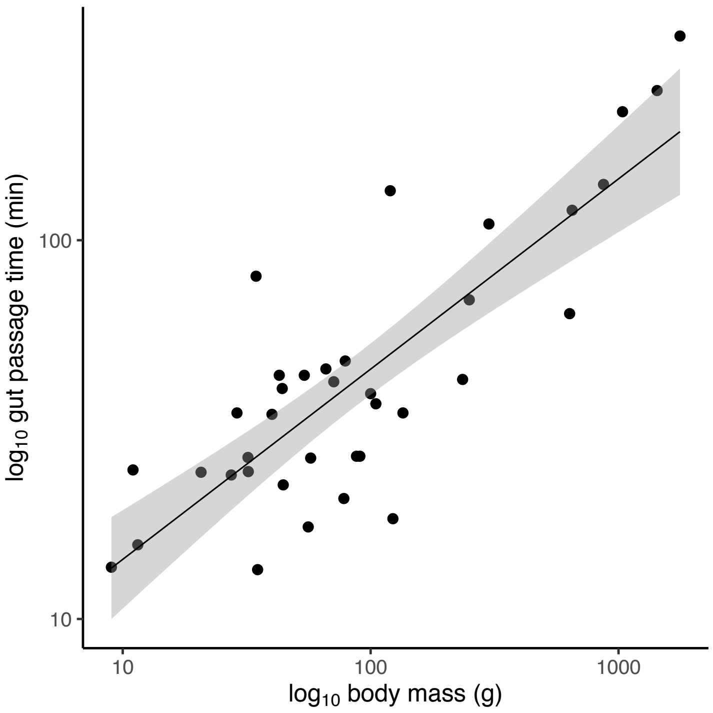

**Fig. A2**. Relationship between **body mass** and **mean dispersal distance** using data extracted from empirical studies of seed dispersal by frugivorous birds (see Table S2 for included studies). Body mass is positively related to mean dispersal distance ($r^2 = 0.4$, $p = 0.007$, n = 16). The grey shaded region indicates the confidence interval for the regression.

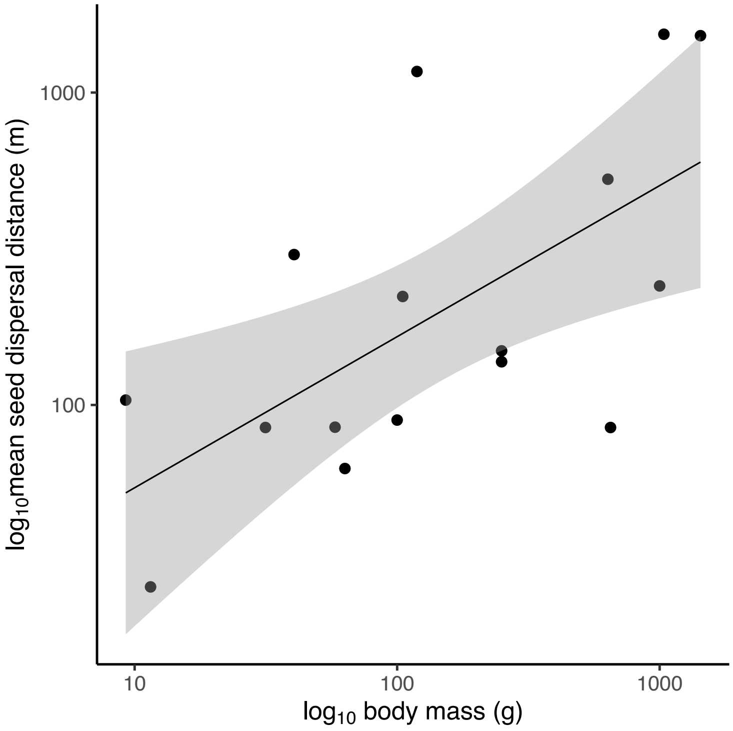

**Fig. A3**. Relationship between **body mass** and **max seed dispersal distance** using data extracted from empirical feeding trials for frugivorous birds (see Table S2). Body mass is positively related to max seed dispersal distance ($r^2 = 0.62$, p = 0.001, n=12). The grey shaded region indicates the confidence interval for the regression.

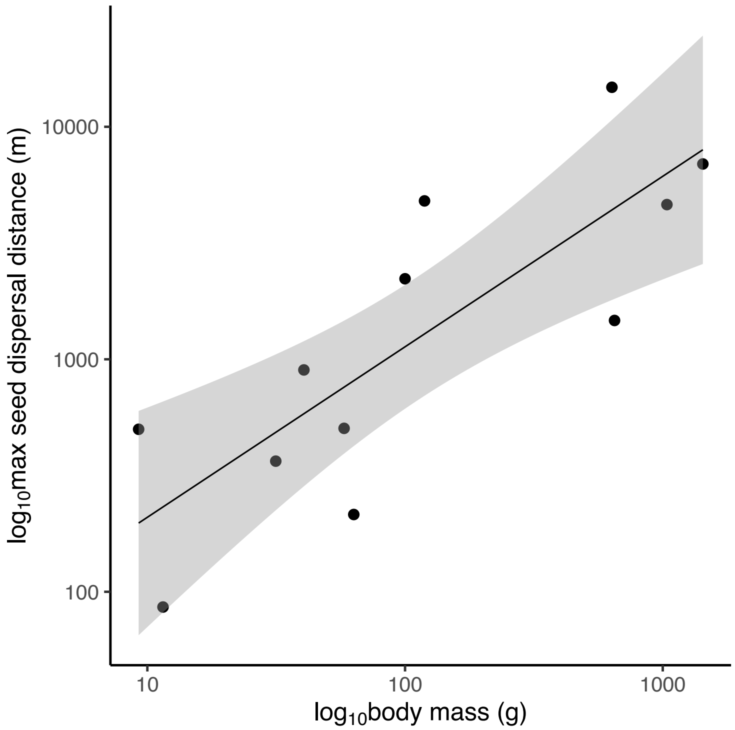

**Appendix 4.**

**Fig. A4.** Long-distance seed dispersal (LDD) results for (b) small, (c) medium, and (d) large fruited plant species.

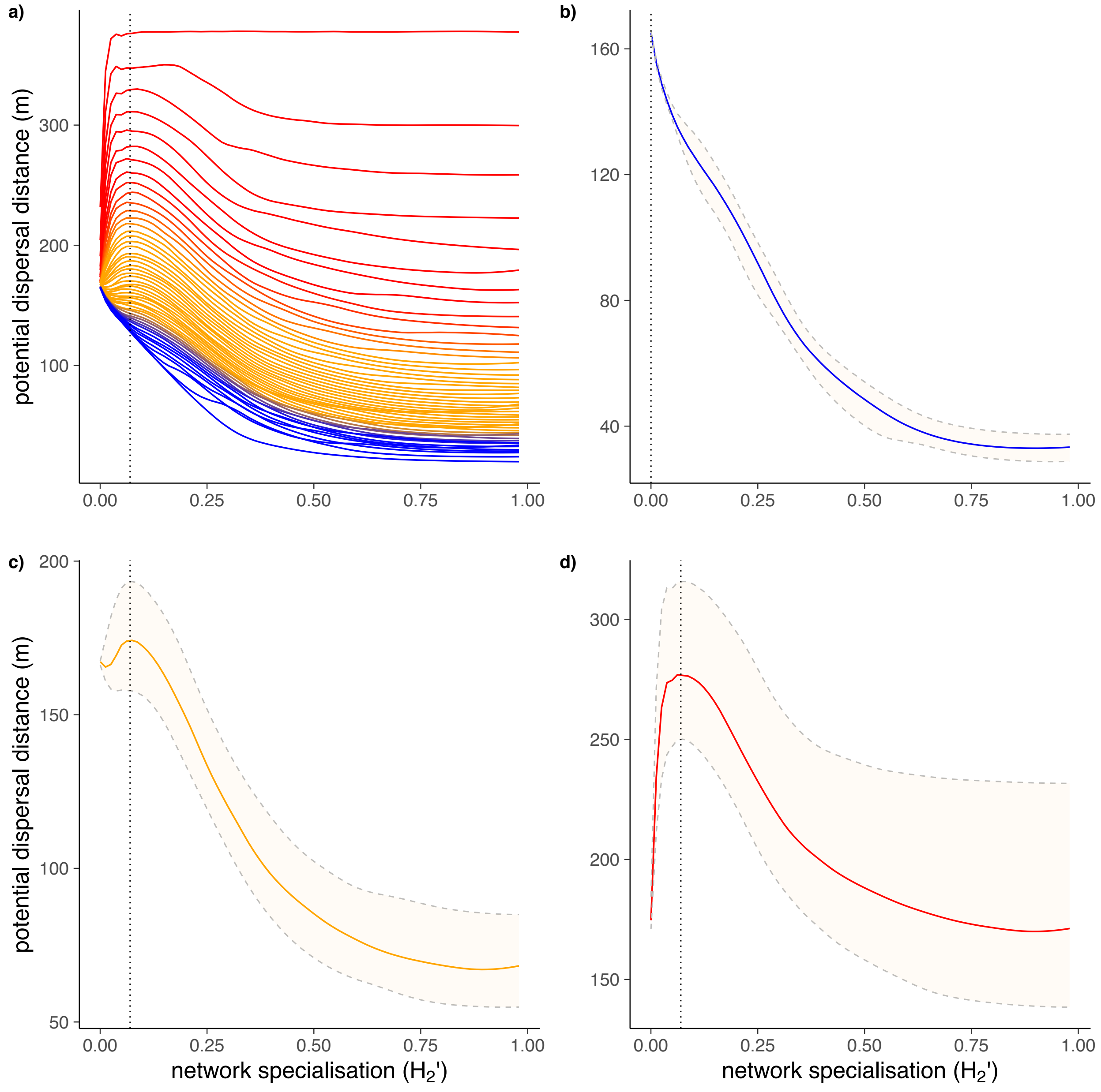

**Appendix 5: Sensitivity analysis**

Morris's elementary effects method estimates the effect of each factor on the model output repeatedly, while the other factors take on different values from their entire ranges, and then averages these estimates into a measure of overall effect; these effects are called elementary effects. The elementary effects are statistically analysed to measure their relative importance (Thiele *et al*. 2004). We used the estimated mean of the distribution of the absolute values of the elementary effects, $\mu^*$, as a sensitivity measure to establish the overall impact of a parameter on the output.

We performed the sensitivity analysis on five model parameters (*k*; Table 1), which were varied according to predefined ranges (see Table S3). The number of tested settings is given by $r \times (k + 1)$, where *r* is the number of elementary effects computed per parameter. As we chose 160 elementary effects, this led to 160 x (5 + 1) = 960 model runs. We ran the global sensitivity analysis for both, the mean and the 95% quantile of seed dispersal distances.

We used the following methods to determine the range of the parameter values to be included in the global sensitivity analysis. For $GPT^{exp}$ we used the 95% confidence intervals of the exponent from the fitted allometric equation; for *GPT*var we used the min and max values from feeding trial studies (Table S1); for $FS^{exp}$ we took the range of 95% confidence intervals of the exponent from those calculated in a similar flight speed allometric equation presented in Alerstam *et al.* 2007; for *FS*sd we took the min and max standard deviation values from those reported from empirical flight speed data in Alerstam *et al*. 2007; for the *CorrFactor* we simply used a min value that was half of the estimated value and a maximum value that was twice the estimated value.

**Table A3.** Sensitivity analysis model parameters and results from the Morris screening method. The top three most influential parameters for median seed dispersal distances are bolded in black; the top three most influential parameters for the 95% quantile of seed dispersal are bolded in orange. μ* is an estimate of the overall influence of a factor on the model output (including interactions with other factors), and σ is an estimate of how much the influence of a factor depended on interactions and stochasticity.

| | | | median | | 95% quantile | |
|---|---|---|---|---|---|---|
| parameter | description | range | $\mu^*$ | σ | $\mu^*$ | σ |
| gut passage time: | | | | | | |
| $GPT^{exp}$ | exponent of the GPT equation (3) | 0.39–0.62 | **0.31** | **0.58** | **0.16** | **0.31** |
| *GPT* var | variance of the GPT gamma distribution, $s^2$ in equation (5) and (6) | 2613–931509 | **1** | **1** | **1** | **0.98** |
| bird movement: | | | | | | |
| $FS^{exp}$ | exponent of the FS equation (4) | 0.13–0.21 | 0.12 | 0.23 | 0.01 | 0.007 |
| *FS* sd | standard deviation of the FS gaussian distribution | 0–4.7 | 0 | 0 | 0 | 0 |
| *CorrFactor* | *fc* in equation (7) | 0.001–0.004 | **0.58** | **0.96** | **0.55** | **1** |

**Fig. A5.** The relationship between network specialisation (*$H_2'$*) and median community seed dispersal distances (TDK$_{community}$) when using the a) min *$GPT^{exp}$* value, and b) max *$GPT^{exp}$*. *$GPT^{exp}$* is included in the top three most influential parameters for: mean and the 95% quantile of seed dispersal distances. Results show the same hump-shaped pattern between *$H_2'$* and community-wide median seed dispersal distances.  Absolute distance values for both the mean (min*$GPT^{exp}$*: peak in seed dispersal = 98 m; max*$GPT^{exp}$*: peak in seed dispersal = 53 m ) and the 95% quantile values (min*$GPT^{exp}$*: peak in seed dispersal = 209 m; max*$GPT^{exp}$*: peak in seed dispersal = 140 m ) are different. Please note different scales of the y-axes.

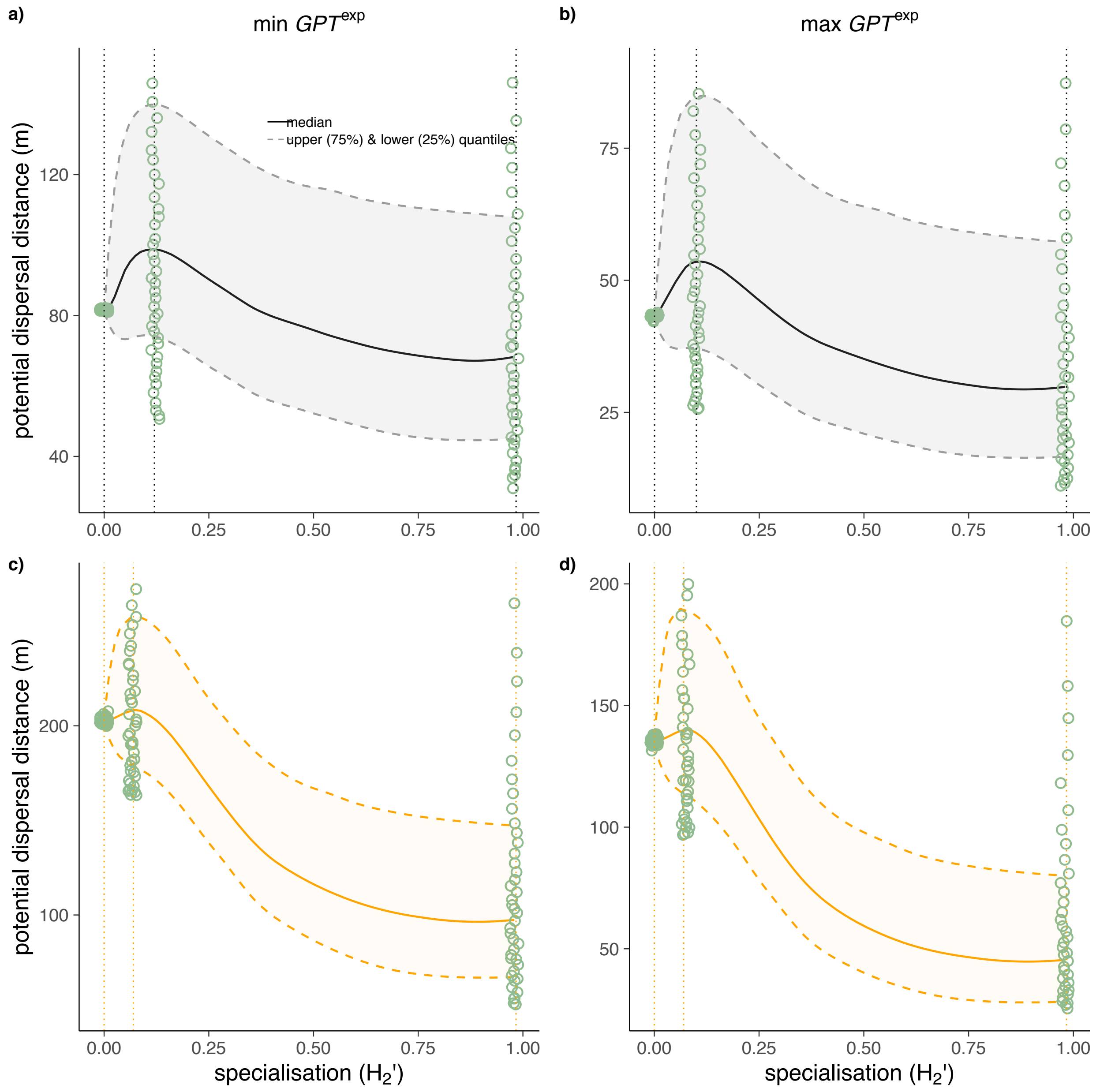

**Fig. A6.** The relationship between network specialisation (*H₂´*) and **mean** community seed dispersal distances ($TDK_{community}$) when using the a) min *GPT*var value, and b) max *GPT*var. *GPT*var is included in the top three most influential parameters for: mean, and 95% quantile seed dispersal distances. All figures show the same hump-shaped pattern between $H_2'$ and mean or LDD community-wide seed dispersal distances. c), and d) report results from the 95% quantile. Absolute seed dispersal distance values were very different for the mean (min*GPT*var: peak in seed dispersal = 2.5 m; max*GPT*var: peak in seed dispersal = 909) and 95% quantile of seed dispersal distances (min*GPT*var: peak in seed dispersal = 6 m; max*GPT*var: peak in seed dispersal = 2157 m).

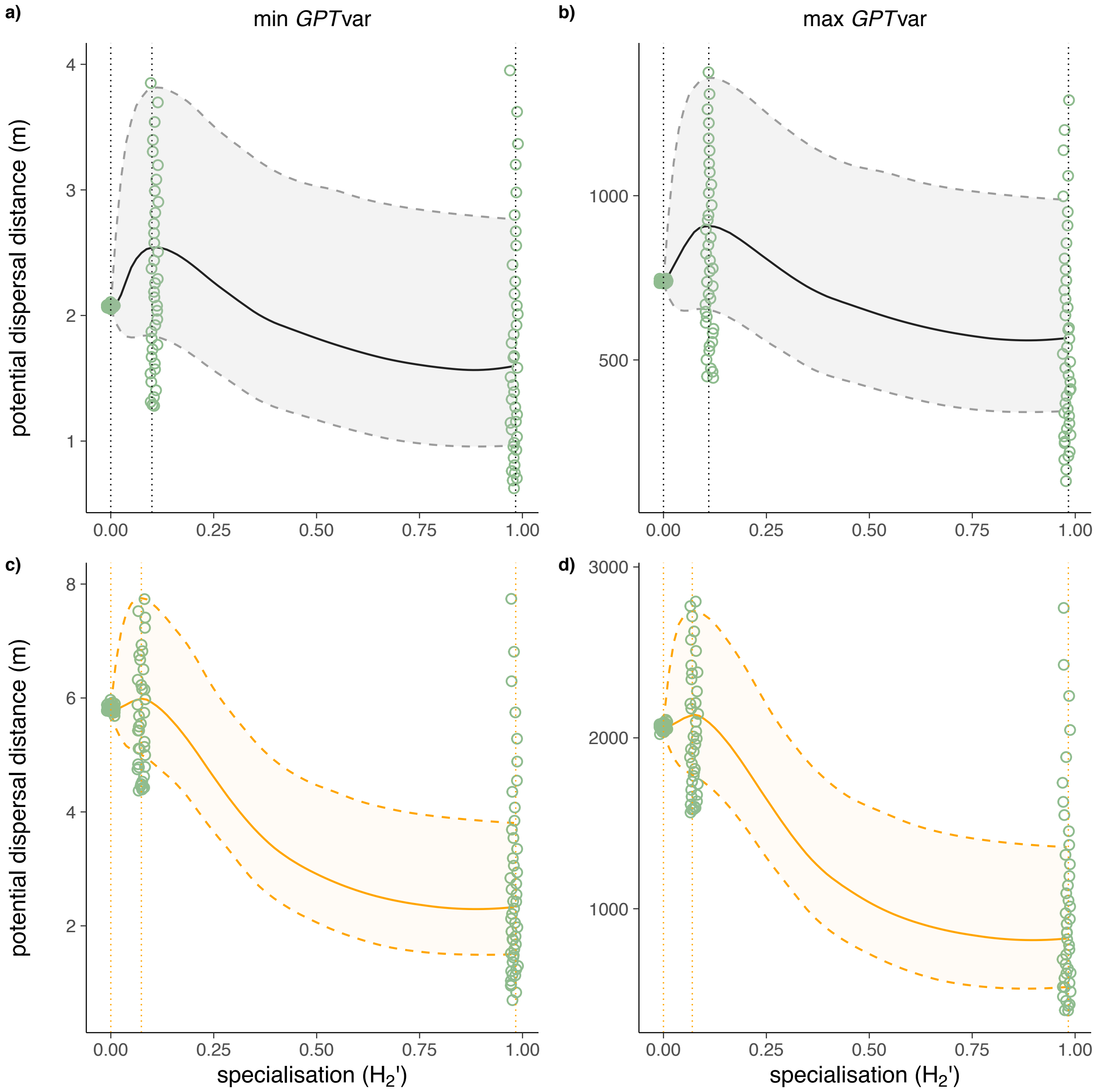

**Fig. A7.** The relationship between network specialisation ($H_2'$) and community seed dispersal distances ($TDK_{community}$) when using the a) min *CorrFactor* value, and b) max *CorrFactor*. *CorrFactor* is included in the top three most influential parameters for: mean, and 95% quantile seed dispersal distances. c), and d) report results from the 95% quantile. All figures show the same hump-shaped pattern between $H_2'$ and median or LDD community-wide seed dispersal distances. Absolute seed dispersal distance values were longer under the max *CorrFactor* scenario. Please note different scales of the y-axes.

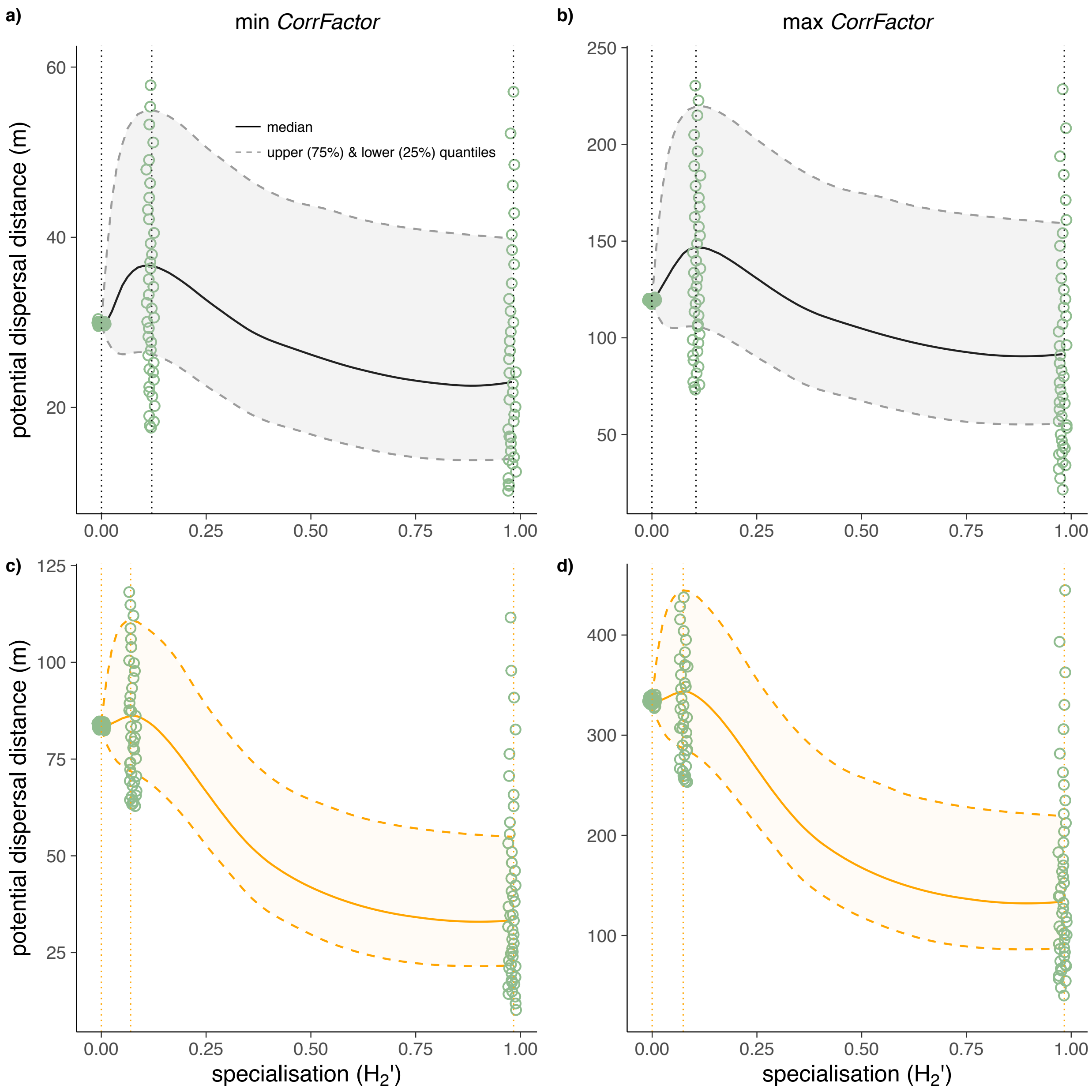
